# Octopaminergic/tyraminergic *Tdc2* neurons regulate biased sperm usage in female *Drosophila melanogaster*

**DOI:** 10.1101/2022.02.14.480434

**Authors:** Dawn S. Chen, Andrew G. Clark, Mariana F. Wolfner

## Abstract

In polyandrous internally fertilizing species, a multiply-mated female can use stored sperm from different males in a biased manner to fertilize her eggs. The female’s ability to assess sperm quality and compatibility is essential for her reproductive success, and represents an important aspect of postcopulatory sexual selection. In *Drosophila melanogaster*, previous studies demonstrated that the female nervous system plays an active role in influencing progeny paternity proportion, and suggested a role for octopaminergic/tyraminergic *Tdc2* neurons in this process. Here, we report that inhibiting *Tdc2* neuronal activity causes females to produce a higher-than-normal proportion of first-male progeny. This difference is not due to differences in sperm storage or release, but instead is attributable to the suppression of second-male sperm usage bias that normally occurs in control females. We further show that a subset of *Tdc2* neurons innervating the female reproductive tract is largely responsible for the progeny proportion phenotype that is observed when *Tdc2* neurons are inhibited globally. On the contrary, overactivation of *Tdc2* neurons does not further affect sperm storage and release or progeny proportion. These results suggest that octopaminergic/tyraminergic signaling allows a multiply-mated female to bias sperm usage, and identify a new role for the female nervous system in postcopulatory sexual selection.

## Introduction

In the fruit fly *Drosophila melanogaster*, females typically mate multiple times with multiple mates throughout their lifetimes (Giardina *et al*. 2017), and store sperm in specialized sperm storage organs (SSO) for days to weeks (reviewed in Neubaum and Wolfner 1999; Bloch Qazi *et al*. 2003). Sperm storage in a multiply-mated female enables postcopulatory sexual selection, where the female can bias sperm usage towards a certain male for fertilization (cryptic female choice), and for different males’ ejaculates to compete for fertilization opportunities (sperm competition) (Parker 1970; Birkhead and Hunter 1990; Eberhard 1996; Birkhead and Møller 1998; Simmons 2002). Both phenomena can interact to give rise to differential paternity proportion, where each of a multiply-mated female’s previous mates enjoy a different paternity proportion. Experiments on differential paternity proportion typically use a double mating paradigm, and the proportion of progeny fathered by the first and second male are referred to as P1 and P2, respectively.

Sperm handling involves many steps in *D. melanogaster*, including sperm mixing, displacement, ejection, storage, and release. *D. melanogaster* females have two types of SSOs: the seminal receptacle (SR) and a pair of spermathecae (Sp). The SR stores the majority of sperm, and the sperm in the SR (but not the Sp) are used for fertilization (i.e. constitutes the “fertilization set”; (Lefevre and Jonsson 1962; Gilbert 1981; Parker 1984; Neubaum and Wolfner 1999; Pitnick *et al*. 1999; Manier *et al*. 2010)). Soon after the start of a second mating, females release a small amount of sperm from the SR into the bursa (Snook and Hosken 2004; Manier *et al*. 2010). The incoming sperm then mix with and displace resident stored sperm until they become well mixed in the bursa and the SR (Manier *et al*. 2010, 2013a). Between 0.5-6 h after mating, females eject excess sperm along with the mating plug from their bursa, thereby terminating sperm mixing and displacement (Manier *et al*. 2010; Lüpold *et al*. 2013; Lee *et al*. 2015). Factors that influence the paternity outcome in wild-type double matings include 1) relatively longer and slower sperm are better at withstanding displacement (as resident sperm) and displacing resident sperm (as incoming sperm); 2) the number of resident and incoming sperm at the time of the second mating; 3) ejection of mating plug before resident and incoming sperm become well mixed, thereby increasing P1 and decreasing P2 (Lüpold *et al*. 2012, 2013). Aside from sperm, several seminal fluid proteins (Sfps) also influence a male’s P1 or P2 (Clark *et al*. 1995; Chapman *et al*. 2000; Fiumera *et al*. 2005, 2007; Wong *et al*. 2008; Fricke *et al*. 2009; Avila *et al*. 2010; Chow *et al*. 2010; Greenspan and Clark 2011; Zhang *et al*. 2013; Castillo and Moyle 2014; Lee et al. 2015; Avila and Wolfner 2017).

Population genetic studies have demonstrated that the female genetic background influences the relative paternity proportions of her mates (Clark and Begun 1998; Clark *et al*. 1999; Chow *et al*. 2010, 2013; Lüpold *et al*. 2013), but the genetic basis of such influence is largely unknown. A GWAS using females from the *Drosophila* Genetic Reference Panel (DGRP) identified 33 genes that harbor variants in mated females most strongly associated with P1, and these genes are enriched for nervous system expression and/or neural function (Chow *et al*. 2013). Further functional tests confirmed that eight candidate genes do indeed affect P1 when knocked down with RNAi in the female’s whole body, nervous system, or subsets of neurons. Of note, knocking down *caupolican* (*caup*), a gene that encodes a transcription factor involved in development, in a female’s octopaminergic/tyraminergic (*Tdc2*) neurons reduces P1 (Chen *et al*. 2019). We hypothesize that *Tdc2* neuron-specific knockdown of *caup* affects the neurons’ development or function, thereby impacting P1. Further investigation was required to establish if and how *Tdc2* neurons, and the biogenic amines octopamine (OA) and tyramine (TA), mediate postcopulatory sexual selection and affect paternity proportion.

OA and TA are synthesized from a tyrosine precursor. Tyrosine decarboxylase (Tdc) first converts tyrosine to TA, and tyramine β-hydroxylase (Tβh) then converts TA to OA (Figure 1A) (Monastirioti *et al*. 1996; Cole *et al*. 2005). *Tdc2* neurons produce OA and TA, and are central to the induction of a suite of physiological and behavioral changes in females after mating, collectively known as postmating responses (PMRs). The PMRs known to be regulated by *Tdc2* neurons include oogenesis, ovulation, egg laying, and refractoriness to remating (Monastirioti *et al*. 1996; Cole *et al*. 2005; Middleton *et al*. 2006; Rodríguez-Valentín *et al*. 2006; Rubinstein and Wolfner 2013; Rezával *et al*. 2014; Meiselman *et al*. 2018; Yoshinari *et al*. 2020). Using singly-mated *Tβh* and *Tdc2* mutant females, Avila et al. showed that OA is also required for sperm release from the SR, and both OA and TA are required for efficient sperm release from both types of SSOs (Avila *et al*. 2012). There are fewer than 200 *Tdc2* neurons in the brain and ventral nerve cord (VNC), innervating the adult central and peripheral nervous systems (Busch *et al*. 2009; Schneider *et al*. 2012; Pauls *et al*. 2018). In particular, a subset of approximately nine *Tdc2* neurons that co-express the sex determination gene *doublesex* (*dsx*) innervate the female RT and mediate PMRs (Rezával *et al*. 2014). These neurons’ cell bodies are located in the abdominal ganglion of the VNC, and they send innervations throughout the female RT (Rezával *et al*. 2014; Yoshinari *et al*. 2020).

**Figure 1.**
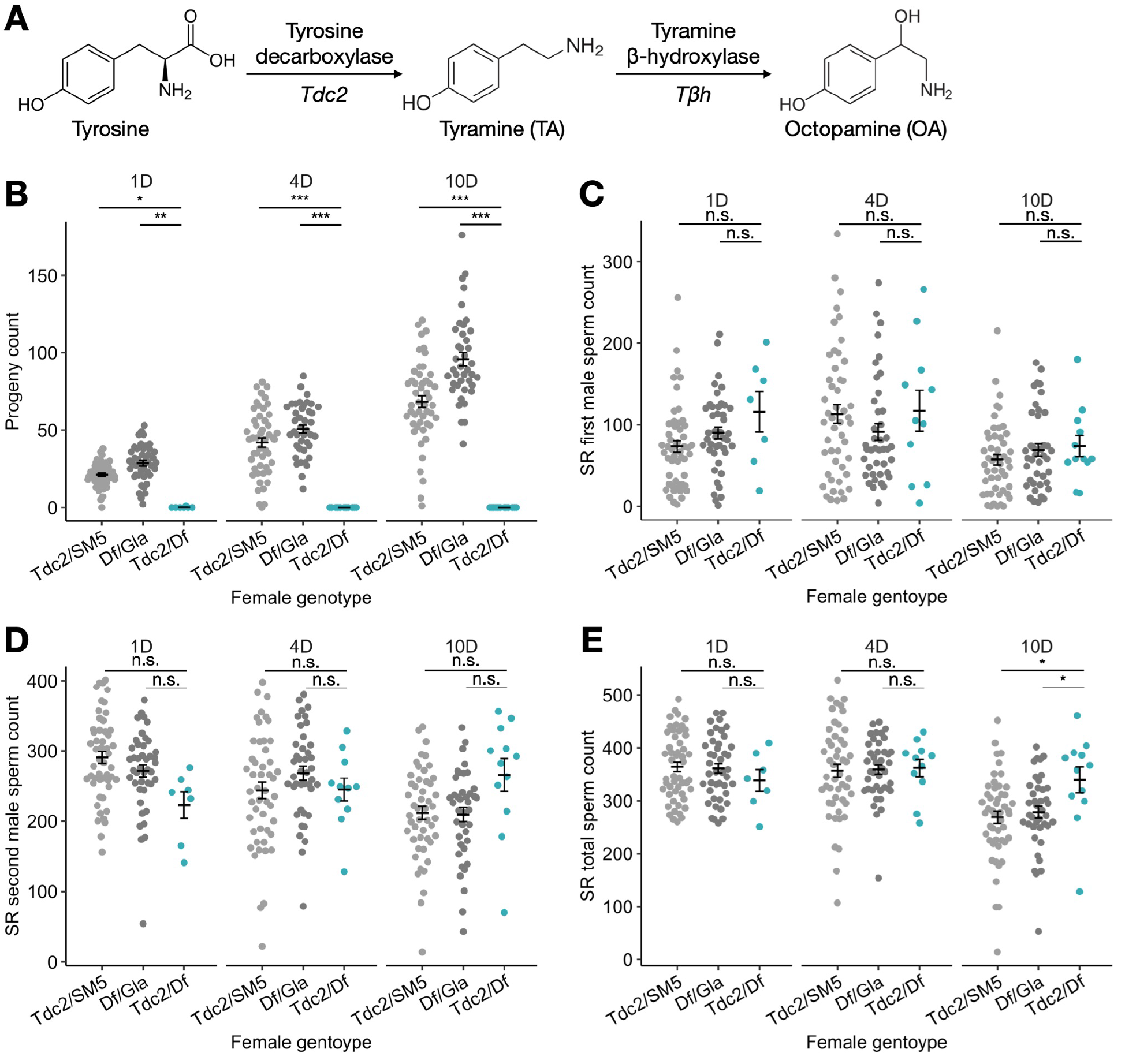
OA/TA-less *Tdc2^RO54^* females are sterile and retain sperm. (A) Biosynthetic pathways of tyramine (TA) and octopamine (OA) from tyrosine, showing the structure of the molecules, the enzymes involved, and the genes in *D. melanogaster* that encode the enzymes. (B-E) *Tdc2^RO54^/SM5* (light gray), *Df(2R)42/Gla* (dark gray), and *Tdc2^RO54^/Df(2R)42* (blue) females mated first to *ProtB-GFP* males then to *ProtB-RFP* males. (B) Progeny count of *Tdc2^RO54^/SM5, Df(2R)42/Gla*, and *Tdc2^RO54^/Df(2R)42* females up to 1, 4, and 10 d AESM. (C-E) Counts of first-male (C), second-male (D), and the sum of first- and second-male sperm (E) in the SR of *Tdc2^RO54^/SM5, Df(2R)42/Gla*, and *Tdc2^RO54^/Df(2R)42* females at 1, 4, and 10 d AESM. Error bars represent mean +/- SEM. Significance levels: * p<0.05, ** p<0.01, *** p<0.001, n.s. not significant.

Because of OA and TA’s involvement in sperm release after a single mating, in this study, we ask if these molecules, as well as *Tdc2* neurons, are also important for differential sperm usage, affecting paternity proportion after two matings. We first report that, similar to singly-mated *Tdc2* mutant females, doubly-mated *Tdc2* mutant females also exhibit sperm retention. We also show that inhibiting *Tdc2* neuronal activity in females increases P1 not by affecting the relative sperm proportion in storage, but by suppressing the females’ biased usage of the second male’s sperm. The P1-increasing effect persists even when the female is first mated to a heterospecific *D. simulans* male. We further show that the *Tdc2/dsx* neurons innervating the female RT are largely responsible for the P1 difference. In contrast, experimentally activating *Tdc2* neurons has limited effects on the progeny paternity proportion or sperm proportion in storage. Our results demonstrate a new mechanism by which the female nervous system can mediate postcopulatory sexual selection.

## Materials and methods

### Fly stocks and husbandry

Fly stocks and crosses were maintained on standard glucose-yeast-agar food on a 12-h light/dark cycle at 22°C, with the exception of *Tdc2>dTrpA1* experiments. In those experiments, all female flies that were assayed developed and aged at 18°C until the start of the experiment (Rezával *et al*. 2014). All flies were collected in single-sex vials soon after eclosion and aged 3-5 days before the start of the experiment.

The stocks we used were Canton S (CS), *Tdc2^RO54^/Gla* (Cole *et al*. 2005), *Df(2R)42/SM5* (BDSC line 3367), *Tβh^nM18^/FM7* (Monastirioti *et al*. 1996), *Tdc2-GAL4* (BDSC line 9313; (Cole *et al*. 2005)), *dsx^FLP^* (gift from the Goodwin Lab; (Rezával *et al*. 2014)), *UAS-TNT* (BDSC line 28838, gift from the Goodwin Lab; (Sweeney *et al*. 1995)), *UAS-dTrpA1* (BDSC line 26263; (Hamada *et al*. 2008)), *UAS>stop>TNT* (gift from the Goodwin Lab), *LHm Ubnls-EGFP ProtB-GFP* (referred to as *ProtB-GFP*; gift from the Pitnick lab; (Manier *et al*. 2010)), *LHm ProtB-DsRed-monomer* (referred to as *ProtB-RFP*; gift from the Pitnick lab; (Manier *et al*. 2010)) and *D. simulans ppk23^-/-^* (gift from the Ruta lab; (Seeholzer *et al*. 2018)).

### Mating latency, copulation duration, and mating plug ejection time

Females were mated to CS males in pairwise matings. For each pair, we recorded the time when both flies were introduced into the same vial (*t_intro_*), copulation start time (*t_start_*), and copulation end time (*t_end_*). Mating latency was calculated as *t_start_* - *t_intro_*, and copulation duration was calculated as *t_end_* - *t_start_*.

Mating plug ejection time (*t_eject_*) was assayed by two methods. 1) Males were removed from vials immediately after mating, and females were flash frozen in liquid nitrogen at 1.5 h or 2 h after the end of mating. Females were later dissected in 1X PBS to examine the presence of mating plugs in the bursae. 2) Immediately after mating, females were transferred to custom-designed mating plug ejection chambers (Hopkins *et al*. 2019) that were then covered with a piece of coverslip and secured with tape on diagonal corners. Mating plug ejection chambers measure 34 mm × 33 mm × 9 mm with a concave depression of 20 mm × 20 mm × 7 mm, and were 3D-printed with black PLA filament at MannUfactory at Cornell University. Because of the autofluorescence of a mating plug component protein PEBme (Ludwig *et al*. 1991; Lung and Wolfner 2001; Lee *et al*. 2015; Avila *et al*. 2015; Cohen and Wolfner 2018), mating plugs are visible under a NightSea GFP flashlight. We scored mating plug ejection every 5-10 min by the presence of plug in the chamber and its loss from the female’s ovipositor. We scored ejection until 6 h after *t_end_*, at which point any unejected females were recorded as 6 h and dissected to confirm the mating plug and sperm mass in bursa. The mating plug ejection time was calculated as *t_eject_* - *t_end_*.

### Long-term sperm storage and usage dynamics

Females were first mated to *ProtB-GFP* males in a pairwise manner. The matings were observed, and males were removed after mating. In the evening of the next day, two *ProtB-RFP* males were added to each female vial and left there overnight to increase the probability of copulation, as females are refractory after the first mating. The next day, males were discarded, and females were each transferred to fresh food vials. We designated this time point as 0 h after the end of the second mating (AESM). In *Tdc2^RO45^/Df(2R)42, Tdc2>TNT*, and *Tdc2/dsx>TNT* experiments, 1/3 of the females of each genotype were flash frozen on each of three timepoints: 1 d, 4 d, and 10 d AESM. In *Tdc2>dTrpA1* experiments, to evaluate baseline sperm storage and release, 1/6 of the females of each genotype were flash frozen at 8 h AESM, and another 1/6 at 24 h AESM. To measure genotype × temperature effects on sperm storage, release, usage and progeny production, half of all remaining females were kept at 22°C and the other half were moved to 29°C for 48 h. All females were flash frozen at 72 h AESM.

F1 progeny were reared to adulthood and counted under an Olympus SZX9 stereomicroscope equipped with a FITC filter. Because the *ProtB-GFP* strain also contained *Ubnls-EGFP*, paternity of the F1 progeny could be ascertained by the presence (first male) or absence (second male) of ubiquitous GFP fluorescence. P1 (proportion of first-male progeny) is calculated as the number of GFP progeny over total progeny a female produces after the second mating.

Female lower RTs were dissected in 1X PBS and imaged with an ECHO Revolve microscope to visualize *ProtB-GFP* (FITC channel; 50 ms) and *ProtB-RFP* (TxRED channel; 1200 ms) sperm heads stored in the SR (10X) and each Sp (20X). Sperm heads were manually counted using the annotation tool in the ECHO Pro app (Echo Laboratories) provided with the Revolve microscope and/or with a custom FIJI plugin (Schindelin *et al*. 2012). The proportion of stored sperm from the first male, S1, was calculated as the number of *ProtB-GFP* sperm over the number of total sperm.

### Sperm transfer and displacement

Females were mated to first males the same way as described above, and were mated to *ProtB-RFP* males in individual vials 48 h after the first mating. Two *ProtB-RFP* males were provided for each female, and mating “trios” were observed. Males were removed immediately after mating, and doubly-mated females were flash frozen 1 h AESM. Females were dissected and imaged as above, with the additional visualization of the mating plug in the DAPI channel (5 ms). The presence and distribution of the mating plug and sperm were compared qualitatively, and the number of *ProtB-GFP* sperm heads were quantified with a custom FIJI plugin. Where necessary, images were stitched together with the “pairwise stitching” FIJI plugin (Preibisch *et al*. 2009).

### Sperm motility

Females were mated as described in the “Sperm transfer and displacement” section and dissected in Grace’s supplemented insect medium (Thermo Fisher) under CO_2_ anesthesia 3-4 h AESM. Their SRs were imaged in the TxRED (600 ms) and FITC (50 ms) channels sequentially, within 3-7 mins of anesthesia, and a 10-second video was obtained in each channel. The videos were then scored manually for the approximate overall proportion of motile sperm into three categories: 0%, 50%, or 100%.

### Conspecific sperm precedence

Females were mated with *D. simulans ppk23^-/-^* males (Seeholzer *et al*. 2018) in mass mating vials with 10 females and 20 males overnight on day 0. The next day (day 1), females were isolated into individual vials and males were discarded. In the evening of day 2, females were each transferred to fresh vials and two *D. melanogaster ProtB-GFP* males were introduced, and the trios were left overnight. In the early afternoon of day 3, males were discarded; we designated this time point as 0 h AESM. At 24 h AESM, each female was transferred to a fresh vial. Every 48 h henceforth, females were transferred to a fresh vial, until 216 h (9 d) AESM when they were discarded. F1 progeny were reared to adulthood and scored as described above. Because only female hybrid progeny were viable, we distinguished between the 3 categories of F1 progeny (*mel/sim* females, *mel/mel* females, *mel/mel* males), and used the female F1 progeny counts to calculate P1.

### Statistical analysis

Statistical analysis was performed in R version 4.0.4 (R Core Team 2021) using the following packages: tidyverse version 1.3.0 (Wickham *et al*. 2019), lme4 version 1.1-26 (Bates *et al*. 2015), lmerTest version 3.1-3 (Kuznetsova *et al*. 2017) and emmeans version 1.5.4 (Lenth 2016). Statistical significance was evaluated using linear models to evaluate the effect of female genotype, and where relevant replicate, time and/or temperature, on a given trait. When models involved repeated measures from the same female, linear mixed models were used with female ID as the random effect. We used ANOVA to evaluate the significance of variables and their interactions, and use emmeans to perform pairwise comparisons on estimated marginal means between genotypes. Proportions such as P1 and S1 were arcsine square root transformed in the linear (mixed) model to stabilize the variance. Throughout the study, only females who produced five or more progeny after the second mating were included in analyses involving P1. Where relevant, nominal p-values were FDR adjusted for multiple testing.

### Data availability

All supplementary figures and the custom FIJI sperm counting script are available on figshare. Raw data, images, and scripts used for data analysis are available upon request.

## Results

### Octopamine and tyramine are required for wildtype sperm storage and usage dynamics

We first asked if doubly-mated *Tdc2^RO54^/Df(2R)42* females exhibited differential sperm usage with respect to each male’s sperm (Cole *et al*. 2005). Females were mated first to *ProtB-GFP* males and subsequently to *ProtB-RFP* males to label the paternity of stored sperm. *Tdc2^RO54^/Df(2R)42* females are sterile (Figure 1B; (Cole *et al*. 2005; Rezával *et al*. 2014)), so we were unable to measure P1 in their matings. Instead, we examined the spatiotemporal dynamics of sperm storage and release to see if it might differ in the absence of Tdc2. We observed that the sperm storage and usage dynamics of each male’s sperm in the SR were similar between *Tdc2^RO54^/Df(2R)42* and control females, with the exception of sperm retention at 10 d AESM (Figure 1C-E). However, in the Sp, *Tdc2^RO54^/Df(2R)42* females stored more first-male sperm at 1 d AESM, and retained more sperm overall at 4 d and 10 d AESM than controls (Figure S1A-C). Combining the sperm counts in both types of SSOs revealed that *Tdc2^RO54^/Df(2R)42* females stored more first-male sperm at 1 d AESM (due to the increased first-male sperm count in the Sp; Figure S1D), retained more second-male sperm at 10 d AESM (Figure S1E), and retained more sperm overall in all SSOs combined by 4 and 10 d AESM (Figure S1F).

These results suggested a potential role for OA and TA in differentially storing or releasing first-versus second-male sperm. We also extended the previous study (Avila *et al*. 2012), showing that both OA and TA are required for efficient sperm release in doubly-mated, as well as singly-mated, females. This result demonstrated that a second mating did not rescue the sperm retention defect, and that the female’s OA and TA signaling are epistatic over male Sfps that promote sperm release from SSOs, such as Sex Peptide (Avila *et al*. 2010).

We also attempted to compare the sperm storage and usage dynamics between doubly-mated *Tβh^nM18^/Tβh^nM18^* mutant and *Tβh^nM18^/FM7* control females (Monastirioti *et al*. 1996). Unfortunately, none of 468 *Tβh^nM18^/Tβh^nM18^* singly-mated females remated with the second male. This strong refractoriness to remating differs from Rezával et al.’s finding that *Tβh^nM18^/Tβh^nM18^* females had abolished PMR and remated more readily than control females did (Rezával *et al*. 2014). Potential sources of differences between our and Rezával et al.’s study could be different male genotypes, mutations that accumulated in the *Tβh^nM18^/FM7* stock over time, or different stock maintenance or assay conditions (the stock in the earlier study was no longer available).

### Inhibiting Tdc2 neurons reduces female fertility and increases P1

To manipulate OA/TA neuronal signaling, we expressed tetanus toxin light chain (*TNT*) in *Tdc2* neurons to inhibit neuronal activity (Sweeney *et al*. 1995). We found that *Tdc2>TNT* females trended towards producing fewer progeny than control females on days 4 and 10 AESM (Figure 2A), but still produced enough progeny for quantification of the relative paternity proportion, P1. When we examined the counts of first- and second-male progeny separately, we found that *Tdc2>TNT* females produced the same amount of first-male progeny as controls, but fewer second-male progeny by day 10 AESM (Figure S2A-B). This progeny count difference also led to *Tdc2>TNT* females’ higher P1 than control females (Fig. 2B). There was no time effect or genotype × time interaction effect on P1 (Figure S2C).

**Figure 2.**
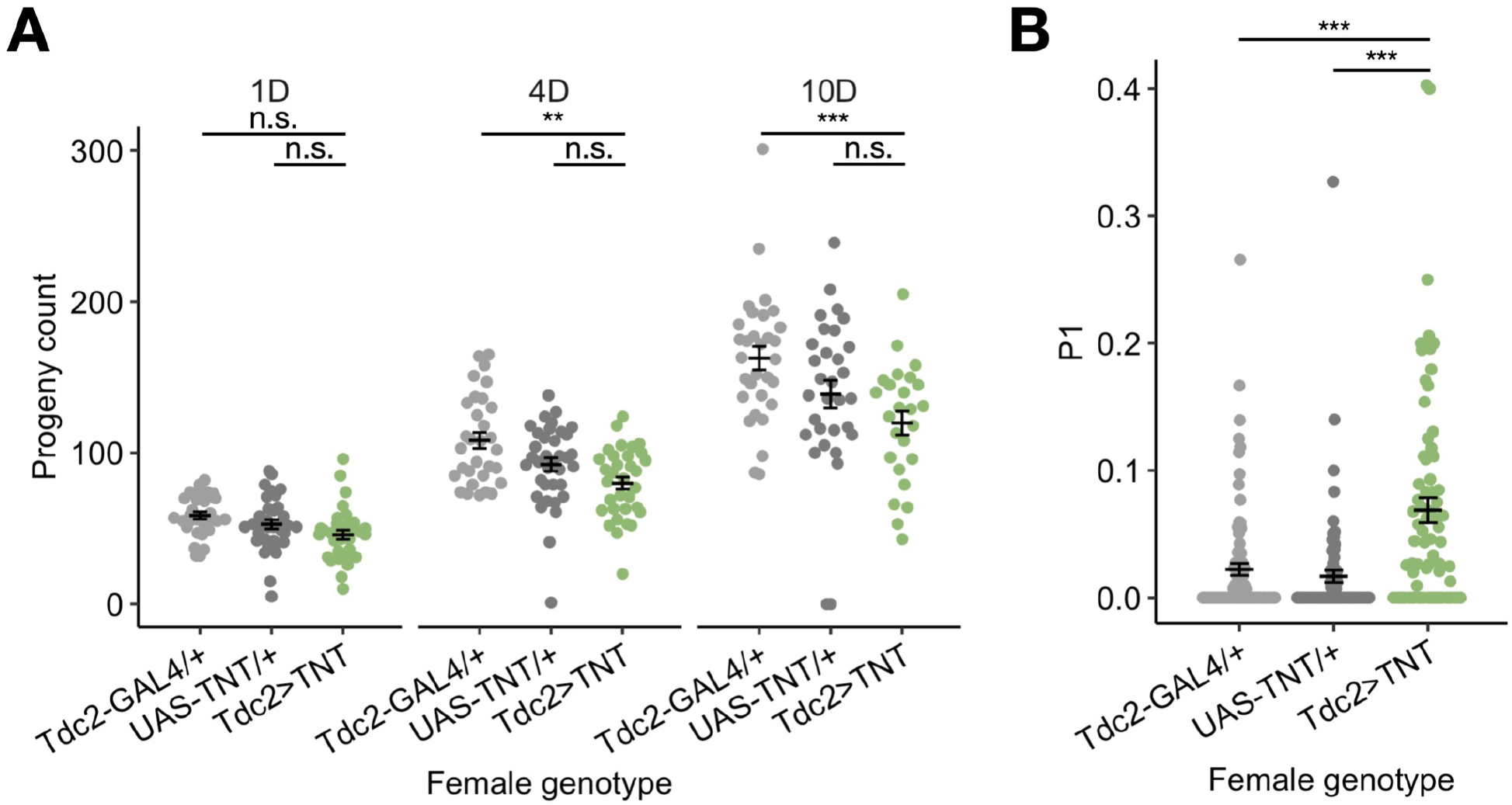
*Tdc2>TNT* females have lower fertility and higher P1. *Tdc2-GAL4/+* (light gray), *UAS-TNT/+* (dark gray), and *Tdc2>TNT* (green) females mated first to *ProtB-GFP* males then to *ProtB-RFP* males. (A) Progeny count of *Tdc2-GAL4/+, UAS-TNT/+*, and *Tdc2>TNT* females up to 1, 4, and 10 d AESM. (B) P1 in *Tdc2-GAL4/+, UAS-TNT/+*, and *Tdc2>TNT* females up to 1, 4, and 10 d AESM. Error bars represent mean +/- SEM. Significance levels: * p<0.05, ** p<0.01, *** p<0.001, n.s. not significant.

To rule out the possibility that the P1 difference was due to female × male genotypic interactions on progeny viability, we mated females of each genotype to either *ProtB-GFP* or *ProtB-RFP* males and calculated the egg-to-adult ratio of their progeny. We did not observe any male genotype effect or female × male genotypic interaction effect on progeny survival (Figure S2D), indicating that the differential paternity outcome is likely due to sperm handling or usage differences, rather than post-zygotic mechanisms.

### Tdc2 neuronal activity is not required for copulation duration or initial sperm handling dynamics

*Tdc2>TNT* females had shorter latency to mating than one of the control genotypes, but similar copulation durations as both control genotypes (Figure 3A-B). We next examined early sperm handling events soon after the second mating, at 1 h AESM. At this time point, the incoming second-male sperm should be actively mixing with and displacing resident first-male sperm, and females have yet to eject the mating plug along with excess unstored sperm (Manier *et al*. 2010). Therefore, we could examine the extent of sperm mixing and displacement, and quantify the number of first-male sperm entering the second mating.

**Figure 3.**
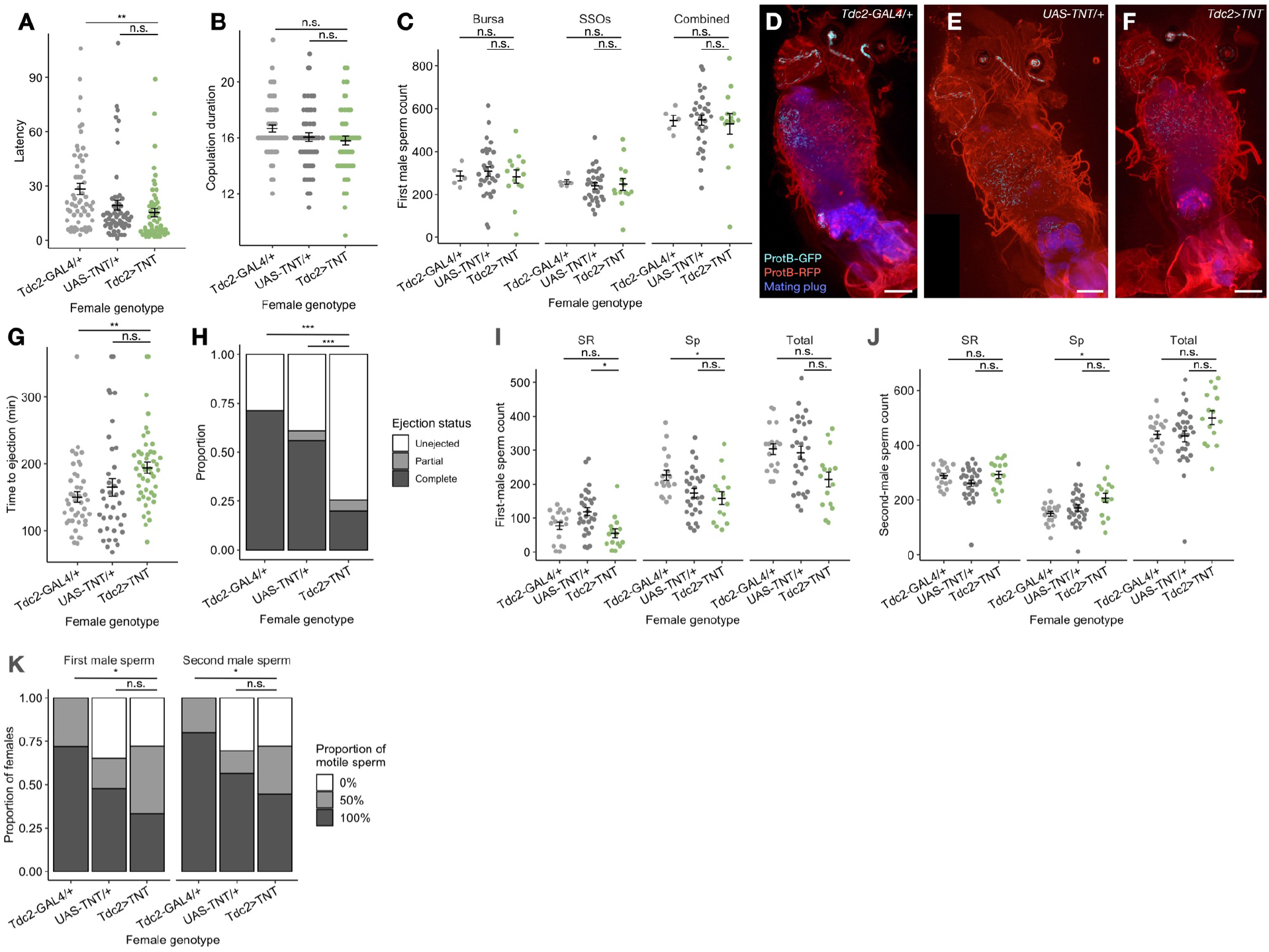
Copulation and sperm handling traits up to 3-4 h AESM are not different between *Tdc2>TNT* and control females. For (A-B, G-H), *Tdc2-GAL4/+* (light gray), *UAS-TNT/+* (dark gray), and *Tdc2>TNT* (green) females mated with CS male. For (C-F, I-K), *Tdc2-GAL4/+* (light gray), *UAS-TNT/+* (dark gray), and *Tdc2>TNT* (green) females mated first to *ProtB-GFP* males then to *ProtB-RFP* males. (A) Latency of *Tdc2-GAL4/+, UAS-TNT/+*, and *Tdc2>TNT* females to copulate. (B) Copulation duration of *Tdc2-GAL4/+, UAS-TNT/+*, and *Tdc2>TNT* females. (C) First-male sperm count of *Tdc2-GAL4/+, UAS-TNT/+*, and *Tdc2>TNT* females in the bursa, SSOs, or the whole lower RT (“Combined”) at 1 h AESM. (D-F) Microscope images of the lower RT in *Tdc2-GAL4/+* (D), *UAS-TNT/+* (E), and *Tdc2>TNT* (F) females, showing first-male sperm heads (teal), second-male sperm heads (red), and mating plug components (blue). The RT structure shows autofluorescence in the TxRED (red) channel. Anterior and posterior regions of the RT are arranged at the top and the bottom, respectively. (G) Mating plug ejection times of *Tdc2-GAL4/+, UAS-TNT/+*, and *Tdc2>TNT* females. Any females who have not ejected their mating plugs by 6 h AESM are recorded as 6 h. (H) Proportions of *Tdc2-GAL4/+, UAS-TNT/+*, and *Tdc2>TNT* females who have not ejected (white), partially ejected (light gray), or completely ejected (dark gray) at 1.5 h AESM. (I-J) Counts of first-male sperm (I) and second-male sperm (J) in the SR, Sp, and both SSOs types combined in *Tdc2-GAL4/+, UAS-TNT/+*, and *Tdc2>TNT* females at 3 h AESM. (K) Proportions of *Tdc2-GAL4/+, UAS-TNT/+*, and *Tdc2>TNT* females with approximately 0% (white), 50% (light gray), or 100% (dark gray) motile first- or second-male sperm in their SR at 3-4 h AESM. Error bars represent mean +/- SEM. Significance levels: * p<0.05, ** p<0.01, *** p<0.001, n.s. not significant. Size bar=100μm.

We found that the number of first-male sperm present in the female RT at the beginning of the second mating was not affected by inhibiting *Tdc2* neuronal activity, and the distribution of first-male sperm among the bursa and the SSOs was also similar across female genotypes (Figure 3C). This result suggested that 1) *Tdc2>TNT* females did not retain more first-male sperm prior to the second mating, and 2) at least at 1 h AESM, *Tdc2>TNT* females were not slower at mixing first- and second-male sperm. The overall sperm distribution throughout the lower female RT was also qualitatively comparable in *Tdc2>TNT* and control females (Figure 3D-F). First- and second-male sperm were well mixed, and did not stratify into clusters. The mating plug also formed normally in *Tdc2>TNT* females (Figure 3D-F).

The timing of mating plug ejection can affect how many first- and second-male sperm are stored, and thereby affect progeny paternity proportion (Lüpold *et al*. 2013; Manier *et al*. 2013b; Lee *et al*. 2015). In general, earlier mating plug ejection after the second mating prevents second-male sperm from sufficiently displacing first-male sperm, and leads to a higher proportion of first-male sperm in the SR (S1) and higher P1 (Manier *et al*. 2010; Lüpold *et al*. 2013). Using two complementary methods to assay mating plug ejection timing, we found that *Tdc2>TNT* females trended towards taking longer to eject the mating plug (Figure 3G), and a larger proportion of *Tdc2>TNT* females were yet to eject at 1.5 h after mating than control females (Figure 3H). Delaying ejection after the sperm are fully mixed should not further affect S1 in the SR, so mating plug ejection is not likely to be responsible for the P1 difference.

We next chose 3 h AESM to examine post-ejection sperm storage, and asked if the number of first- and second-male sperm stored after ejection could contribute to P1 differences. Overall, the sperm storage pattern was comparable across genotypes (Figure 3I-J), consistent with the previous report that OA does not regulate initial sperm entry into storage (Avila *et al*. 2012). *Tdc2>TNT* females stored the same amount of second-male sperm as controls (Figure 3J), and slightly fewer first-male sperm in the SR and Sp than one or the other control genotype, but comparable amounts of first-male sperm overall (Figure 3I). The proportions of motile first- and second-male sperm were also similar across genotypes (Figure 3K). Taken together, these results suggested that *Tdc2* neuron activity was not required for early sperm handling or sperm motility within the SR. Despite the observation that *Tdc2>TNT* females had slightly delayed mating plug ejection, this defect was not likely to explain the higher P1.

### Biased sperm usage explains P1 difference

We next directed our attention to the spatiotemporal dynamics of sperm release and usage between 1 d and 10 d AESM in *Tdc2>TNT* and control females. Female genotype had little effect on the numbers of first- and second-male sperm in the SR, except for a slight trend in *Tdc2>TNT* females towards retaining second-male sperm and a corresponding increase in total sperm count in the SR at day 10 AESM (Figure 4A-C). This small difference did not result in a reduction of S1 in the SR (Figure 4D). In *Tdc2>TNT* females, Sp stored more first-male sperm on day 1 AESM and continued to have more first-male sperm throughout the 10-day assay, which also contributed to *Tdc2>TNT* females having more first- and second-male sperm combined in the Sp than controls (Figure S3A). When combining SR and Sp sperm counts, the overall sperm storage and release pattern is largely consistent across female genotypes, with the most notable difference being a strong sperm retention phenotype at day 10 AESM (Figure S3D-F).

**Figure 4.**
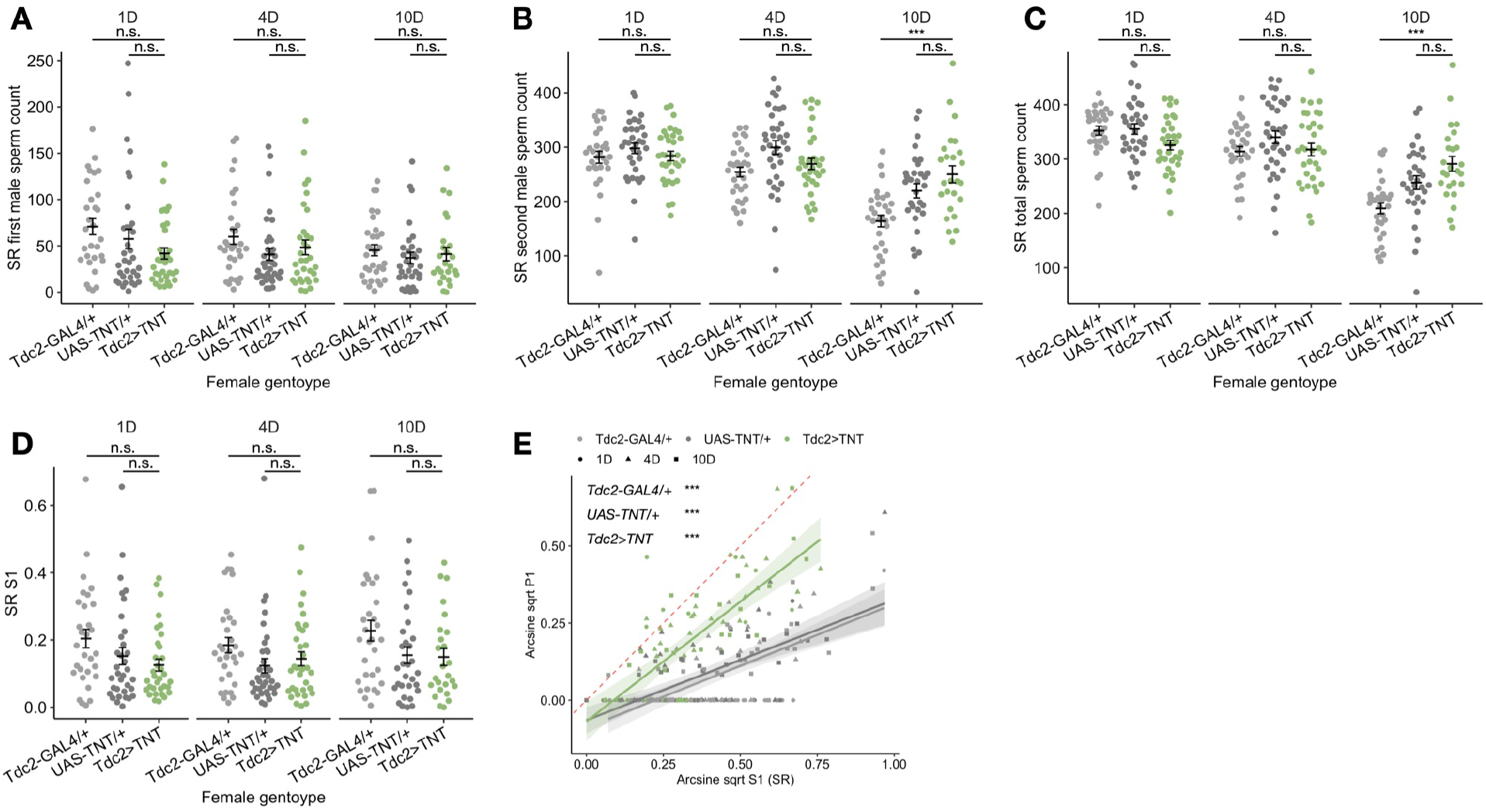
*Tdc2>TNT* females do not store or release more first-male sperm, but suppress sperm usage bias toward second-male sperm. *Tdc2-GAL4/+* (light gray), *UAS-TNT/+* (dark gray), and *Tdc2>TNT* (green) females mated first to *ProtB-GFP* males then to *ProtB-RFP* males. (A-C) Counts of first-male (A), second-male (B), and the sum of first- and second-male sperm (C) in the SR of *Tdc2-GAL4/+, UAS-TNT/+*, and *Tdc2>TNT* females at 1, 4, and 10 d AESM. (D) S1 of *Tdc2-GAL4/+, UAS-TNT/+*, and *Tdc2>TNT* females at 1, 4, and 10 d AESM. (E) Arcsine square root transformed S1 in the SR is correlated with arcsine square root transformed P1. Red dotted line represents the y=x diagonal. Each data point represents one female, with colors corresponding to genotypes and shapes corresponding to time points. Shaded areas around regression lines represent 95% confidence intervals. Significance levels show the strength of correlation between transformed P1 and S1 (SR) for each genotype. Error bars represent mean +/- SEM. Significance levels: * p<0.05, ** p<0.01, *** p<0.001, n.s. not significant.

We examined the correlation between P1 and S1 (after arcsine-square root transformation) to understand how sperm proportion translates to progeny proportion. The slope of the regression line between transformed P1 and S1 can reveal whether females use sperm from either male in a biased manner to produce progeny. If P1 is significantly greater than S1, it suggests that females bias sperm usage toward the first male; and the reverse is true if P1 is significantly lower than S1. All three genotypes we assayed exhibited bias for using second-male sperm to produce progeny (P1<S1), but such bias was stronger in *Tdc2-GAL4/+* and *UAS-TNT/+* control females than in *Tdc2>TNT* females, as shown by their lower slopes (Figure 4E). This result suggested that *Tdc2* neuron activity in control females promoted bias for second-male sperm to be used for progeny production, and inhibiting it allowed females to use sperm from the two males closer to their proportions in the SR.

### Reproductive tract Tdc2/dsx neurons are largely responsible for P1 effects

The small population of *Tdc2/dsx* neurons innervates the female RT and regulates various aspects of the PMR (Rezával *et al*. 2014; Yoshinari *et al*. 2020). We asked if they might also be the subset of *Tdc2* neurons that underlie the P1 and biased sperm usage phenotypes we observed. We used an intersectional strategy combining *Tdc2-GAL4* with *dsx^FLP^* to drive a version of *UAS-TNT* with an FRT-flanked stop cassette inserted upstream of *TNT* (*UAS>stop>TNT*; (Bohm *et al*. 2010)). FLP recombinase excises the stop codon between *UAS* and *TNT* in *dsx^+^* cells to generate functional *UAS-TNT* alleles; if these cells are also *Tdc2^+^, GAL4* drives expression of *TNT* to specifically inhibit these *Tdc2/dsx* neurons.

*Tdc2-GAL4/UAS>stop>TNT; dsx^FLP^/+* (henceforth *Tdc2/dsx>TNT*) females phenocopied *Tdc2>TNT* with respect to fertility (Figure 5A, Figure S4A-B) and P1 (Figure 5B), suggesting that both traits might be mediated by this subset of *Tdc2* neurons. Interestingly, *Tdc2/dsx>TNT* females had a more severe fertility defect than *Tdc2>TNT* females (Figure 5A). We only included females who had produced five or more progeny after the second mating when analyzing P1 because lower fertility affects the “resolution” of P1. *Tdc2/dsx>TNT* females had higher P1 than controls, and we did not observe a time or genotype × time effect on P1 (Figure S4C). The overall spatiotemporal dynamics of sperm storage and release in the SR was similar in *Tdc2/dsx>TNT* females to that in controls, with slightly more rapid release of second-male sperm than one or the other control at day 4 and 10 AESM, and slightly fewer overall sperm stored at the same time points (Figure 5C-E). The sperm loss observation in *Tdc2/dsx>TNT* females is opposite to the sperm retention defect in *Tdc2>TNT* females, which might suggest that non-RT *Tdc2* neurons are responsible for (or can compensate for the loss of RT *Tdc2* neuron activity on) sperm release from storage. Sperm storage and release in the Sp and both SSOs combined were very consistent across genotypes (Figure S4D-I). S1 in the SR was also not significantly different when *Tdc2/dsx* neurons were inhibited (Figure 5F).

**Figure 5.**
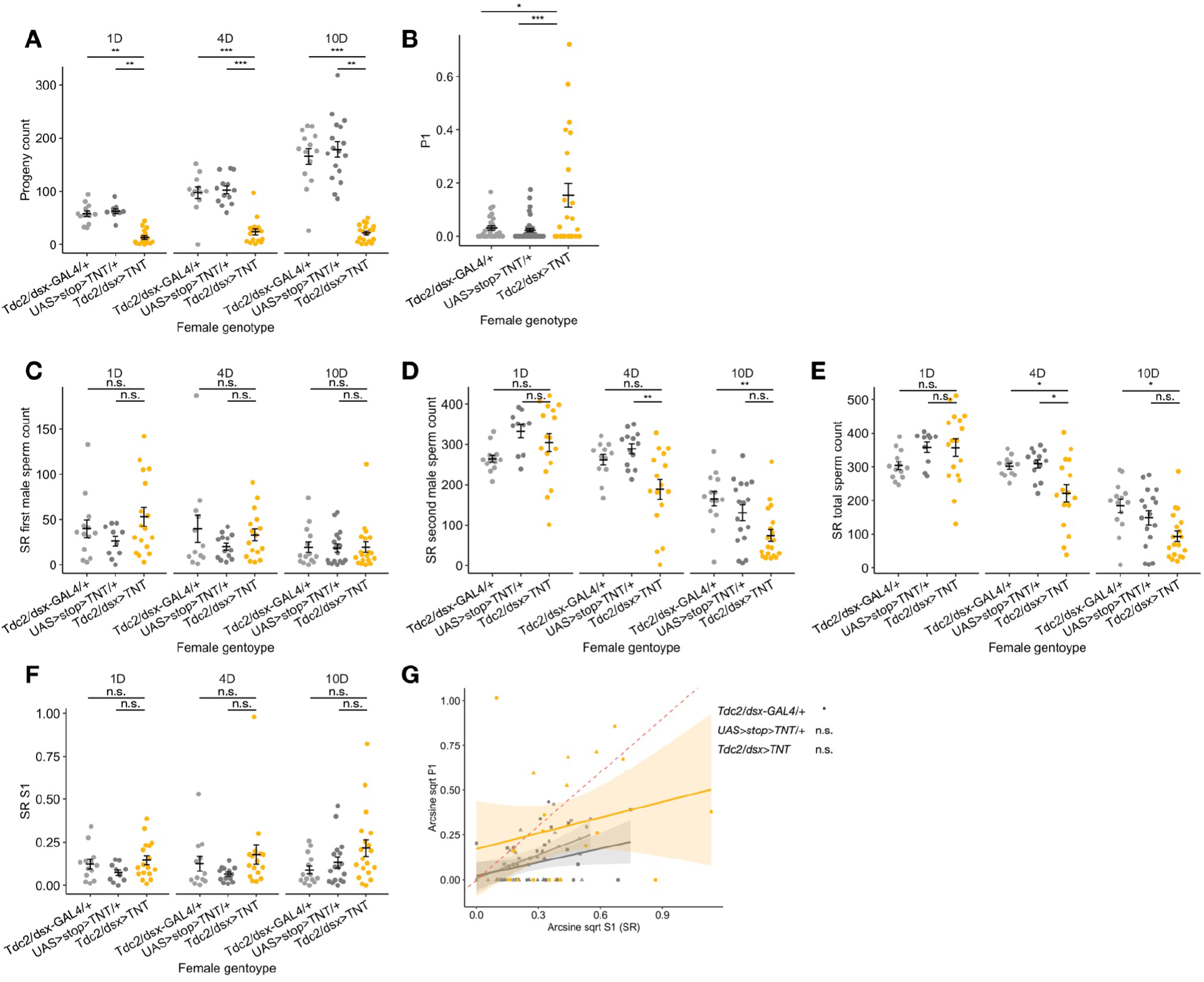
*Tdc2/dsx>TNT* females have lower fertility, higher P1, and do not store or release more first-male sperm. *Tdc2/dsx-GAL4/+* (light gray), *UAS>stop>TNT/+* (dark gray), and *Tdc2/dsx>TNT* (orange) females mated first to *ProtB-GFP* males then to *ProtB-RFP* males. (A) Progeny count of *Tdc2/dsx-GAL4/+, UAS>stop>TNT/+*, and *Tdc2/dsx>TNT* females up to 1, 4, and 10 d AESM. (B) P1 of *Tdc2/dsx-GAL4/+, UAS>stop>TNT/+*, and *Tdc2/dsx>TNT* females up to 1, 4, and 10 d AESM. (C-E) Counts of first-male (C), second-male (D), and the sum of first- and second-male sperm (E) in the SR of *Tdc2/dsx-GAL4/+, UAS>stop>TNT/+*, and *Tdc2/dsx>TNT* females at 1, 4, and 10 d AESM. (F) S1 of *Tdc2/dsx-GAL4/+, UAS>stop>TNT/+*, and *Tdc2/dsx>TNT* females at 1, 4, and 10 d AESM. (G) Arcsine square root transformed S1 in the SR plotted against arcsine square root transformed P1. Red dotted line represents the y=x diagonal. Each data point represents one female, with colors corresponding to genotypes and shapes corresponding to time points. Shaded areas around regression lines represent 95% confidence intervals. Significance levels show the strength of correlation between transformed P1 and S1 (SR) for each genotype. Error bars represent mean +/- SEM. Significance levels: * p<0.05, ** p<0.01, *** p<0.001, n.s. not significant.

Unfortunately, because requiring progeny count greater than or equal to five disproportionately reduced the sample size of *Tdc2/dsx>TNT* females with meaningful P1s, we were unable to determine whether *Tdc2/dsx>TNT* females similarly exhibited reduced biased usage toward second-male sperm (Figure 5G). Nonetheless, since sperm storage and release was at control-like levels in *Tdc2/dsx>TNT* females, it would be likely that they also achieved P1 by altering sperm usage for fertilization, such as suppressing bias toward second-male sperm.

### Tdc2 neurons are required for conspecific sperm precedence

So far we have shown that *Tdc2* neurons mediate biased sperm usage based on the mating order of two *D. melanogaster* males. We further asked if *Tdc2>TNT* females mated first to less compatible males would avoid using first-male sperm, and have control-like levels of lower P1. We mated *Tdc2>TNT* and control females first with a heterospecific, *D. simulans* male, followed by a second mating with a conspecific, *D. melanogaster ProtB-GFP* male. We used *ppk23^-/-^ D. simulans males*, who were unable to detect the *D. melanogaster* female pheromone and therefore courted *D. melanogaster* females rigorously (Seeholzer *et al*. 2018), to facilitate the first, heterospecific mating.

Similar to the case in conspecific matings, *Tdc2>TNT* females had lower fertility than controls, and particularly fewer progeny were fathered by the second male (Figure 6A). To our surprise, *Tdc2>TNT* females also had higher P1 (Figure 6B), especially at 1-3 d AESM (Figure 6C). This might suggest that the lack of *Tdc2* neuronal activity prevents females from efficiently utilizing the second male’s sperm (as in conspecific double mating experiments) or distinguishing between con- and heterospecific sperm soon after the second mating. Overall, we find that *Tdc2>TNT* female’s suppression of second-male sperm usage bias extends to heterospecific matings, at least shortly after the second mating.

**Figure 6.**
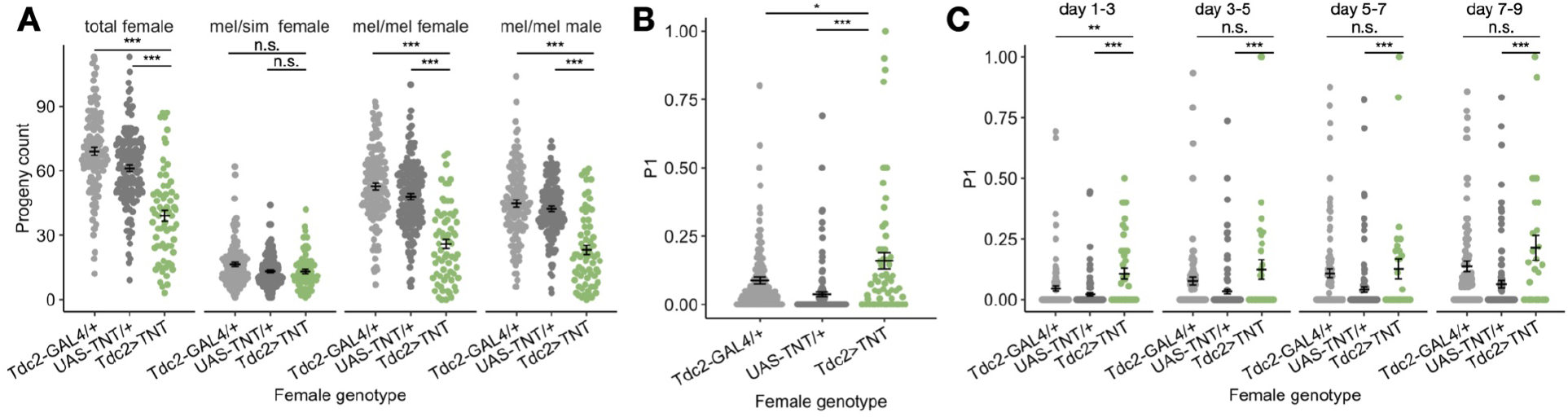
*Tdc2>TNT* females have lower fertility and higher P1 when first mated to heterospecific *D. simulans* male. *Tdc2-GAL4/+* (light gray), *UAS-TNT/+* (dark gray), and *Tdc2>TNT* (green) females mated first to *D. simulans ppk23^-/-^* males then to *D. melanogaster ProtB-GFP* males. (A) Total female progeny, mel/sim hybrid female progeny, mel/mel female progeny, and mel/mel male progeny count of *Tdc2-GAL4/+, UAS-TNT/+*, and *Tdc2>TNT* females between 1 and 9 d AESM. (B) P1 of *Tdc2-GAL4/+, UAS-TNT/+*, and *Tdc2>TNT* females between 1 and 9 d AESM. (C) P1 of *Tdc2-GAL4/+, UAS-TNT/+*, and *Tdc2>TNT* females broken down by ranges of days AESM. Error bars represent mean +/- SEM. Significance levels: * p<0.05, ** p<0.01, *** p<0.001, n.s. not significant.

### Activation of Tdc2 neuronal activity has limited effect on sperm handling

After consistently observing that inhibiting *Tdc2* neuron activity increased the paternity proportion of the first male, we asked if activating *Tdc2* neurons might have the opposite effect. We used the heat-sensitive cation channel *dTrpA1* under UAS control (Hamada *et al*. 2008) to thermogenetically activate *Tdc2* neurons by exposing *Tdc2>dTrpA1* females to 29°C for 48 h, between 24 h and 72 h AESM. We used genotype controls and 22°C temperature controls to assess any genotype × temperature effects on fertility, P1, and the dynamics of sperm storage, release, and usage. We also examined the fertility and sperm storage of females before undergoing temperature treatment, at 8 h and 24 h AESM. We did not find significant genotype effects on fertility, or initial sperm storage or release (Figure 7A, D-F, Figure S5A-F).

**Figure 7.**
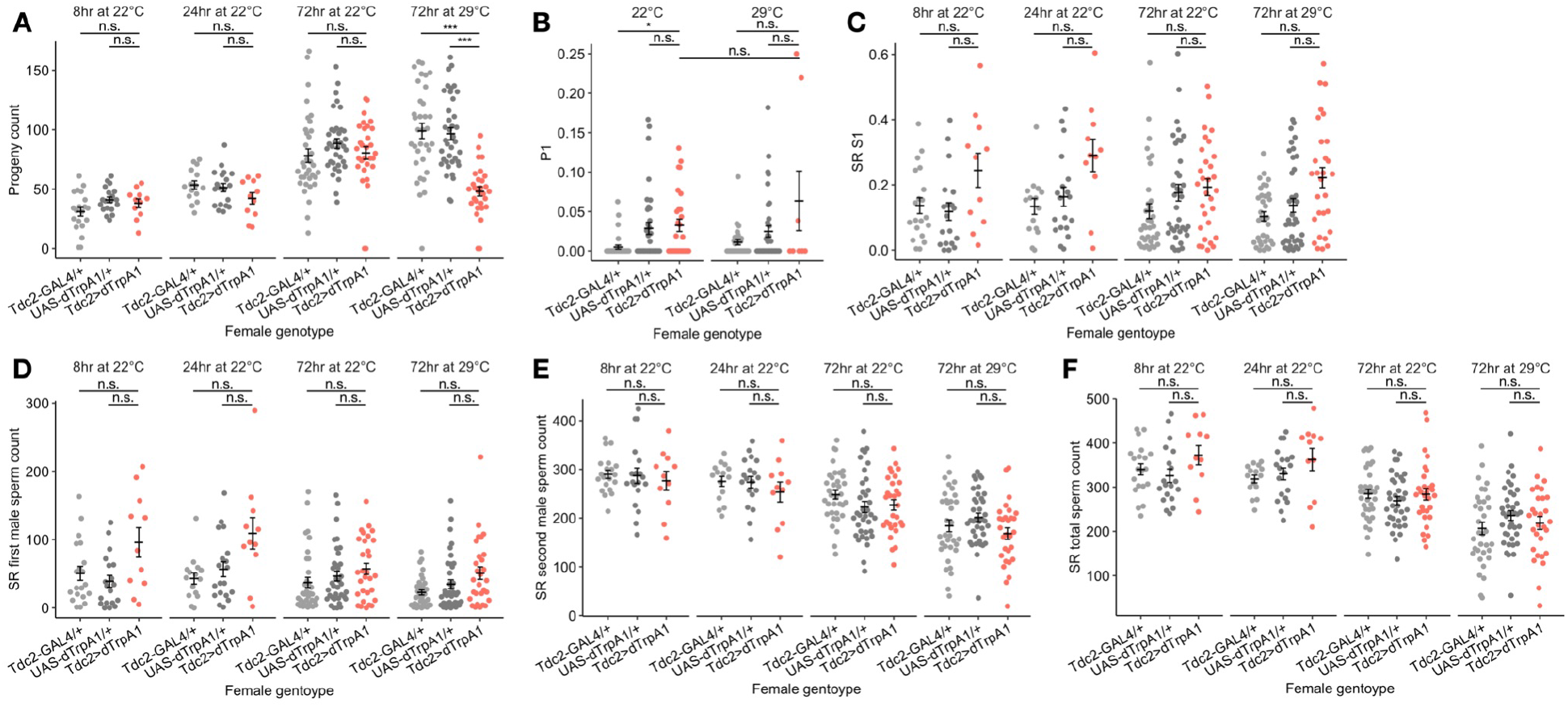
Activating *Tdc2* neurons reduces fertility, but has limited effects on P1 and sperm storage and release. *Tdc2-GAL4/+* (light gray), *UAS-dTrpA1/+* (dark gray), and *Tdc2>dTrpA1* (red) females mated first to *ProtB-GFP* males then to *ProtB-RFP* males. (A) Progeny count of *Tdc2-GAL4/+, UAS-dTrpA1/+*, and *Tdc2>dTrpA1* females up to 8 h, 24 h, and 72 h AESM at 22°C or 29°C. (B) P1 of *Tdc2-GAL4/+, UAS-dTrpA1/+*, and *Tdc2>dTrpA1* females at 22°C or 29°C between 24 h and 72 h AESM. (C) S1 of *Tdc2-GAL4/+, UAS-dTrpA1/+*, and *Tdc2>dTrpA1* females at 8 h, 24 h, and 72 h AESM at 22°C or 29°C. (D-F) Counts of first-male (D), second-male (E), and the sum of first- and second-male sperm (F) in the SR of *Tdc2-GAL4/+, UAS-dTrpA1/+*, and *Tdc2>dTrpA1* females at 8 h, 24 h, and 72 h AESM at 22°C or 29°C. Error bars represent mean +/- SEM. Significance levels: * p<0.05, ** p<0.01, *** p<0.001, n.s. not significant.

Experimentally activating *Tdc2* neurons for 48 h significantly reduced female fertility (Figure 7A). This was not due to the temperature alone, as control females had higher fertility at 29°C (Figure 7A). The lower fertility was not linked to sperm retention, as the high temperature treatment did not affect sperm counts in the SR, Sp or overall sperm counts (Figure 7D-F, Figure S5A-F). P1 and S1 were not affected by *Tdc2* neuron activation (Figure 7B-C).

## Discussion

Reproductive success requires cooperation between females and males in different aspects of reproduction, from recognition, courtship, and copulation, to postmating gamete usage. At the same time, the sexes’ divergent strategies to achieve optimal reproductive output results in sexual conflict. In polyandrous species, complex interactions between the female RT and ejaculate components from previous mates mediate postcopulatory sexual selection and creates differential paternity proportion. Specific female genes have been found to influence paternity proportion (Chow *et al*. 2010; Sitnik *et al*. 2014; Smith *et al*. 2017; Chen *et al*. 2019), but a mechanistic view of how these genes are involved in postcopulatory sexual selection remain largely elusive.

Here, we focus on octopaminergic/tyraminergic *Tdc2* neurons, which have extensive roles in female reproduction and mediate PMRs, and examine how their activity regulates different aspects of sperm handling and progeny production to influence paternity proportion. We found that control females tended to use second-male sperm in a biased manner for producing progeny, which was reflected in their lower P1 values. Inhibiting *Tdc2* neurons, including the *Tdc2/dsx* neurons innervating the female RT, resulted in the loss of this ability. Instead, *Tdc2>TNT* and *Tdc2/dsx>TNT* females used sperm at a proportion more closely matching S1 in the SR, giving rise to generally higher P1. We also examined postzygotic progeny survival and other aspects of sperm handling, including sperm mixing, displacement, ejection, storage, and release, but none of these aspects could explain the P1 difference. In contrast, when we activated *Tdc2* neurons instead, we did not observe an even stronger usage bias towards second-male sperm. This result suggests a “ceiling effect” on the amount of OA in mated females, as has been observed in the context of ovulation (Rubinstein and Wolfner 2013), and/or on the already small number of first-male progeny produced by control females.

The biased sperm usage observed in this study is likely based on mating order, as both tester male strains are in the LHm background (Manier *et al*. 2010) so female × male × male genotypic interactions on sperm handling and usage should be minimal. Studies show that females become more selective of future mates after mating (Jennions and Petrie 2000; Kokko and Mappes 2005; Kohlmeier *et al*. 2021). If the selectiveness is predictive of higher male quality, then it may be beneficial for females to default to preferential use of the last male’s sperm.

We did not find evidence that *D. melanogaster* females stored each male’s sperm in separate storage organs, a pattern that had been observed in *D. simulans* females (Manier *et al*. 2013b). Two potential mechanisms may explain how the female RT biases sperm usage towards the second male for fertilization. 1) Compared to the first male’s sperm, each subsequent male’s sperm encounters a female RT that has already transitioned to a mated state. The transcriptome, neuromodulator, secretion, and muscular contraction states of the mated female RT (McGraw *et al*. 2004, 2008; Middleton *et al*. 2006; Mack *et al*. 2006; Rodríguez-Valentín *et al*. 2006; Adams and Wolfner 2007; Schnakenberg *et al*. 2011; Rubinstein and Wolfner 2013; Heifetz *et al*. 2014; Mattei *et al*. 2015; Newell *et al*. 2020; White *et al*. 2021) may be more efficient at mediating interactions with the incoming ejaculate and better nurture the incoming sperm. The loss or reduction of OA changes the molecular identity and muscle contraction of the female RT (Rodríguez-Valentín *et al*. 2006; Rubinstein and Wolfner 2013; Heifetz *et al*. 2014), and may affect how it interacts with sperm and other ejaculate components. 2) A recent study demonstrates that during sperm storage, sperm progressively lose Sfps and gain female-derived proteins: whereas uniquely female-derived proteins only constitute 1.2% of the sperm proteome at 30 min after mating, this proportion increases to 19.1% by 4 days after mating (McCullough et al. 2022). The relatively more male-like proteome of second-male sperm may allow the female to distinguish those from first-male sperm, or even offer them an advantage to reach the fertilization site or to fertilize the egg. In wild-type females, a few sperm are released from the SR at around the same time when an egg is released into the bursa (Bloch Qazi *et al*. 2003). *Tdc2^RO54^* mutant females and *Tdc2>TNT* females release sperm inefficiently. If they release fewer sperm at a time, sperm may be used for fertilization in closer proportion to their representation in the SR. Advances in long-term live imaging technologies may allow future studies to reveal how sperm release is coordinated with ovulation, and how the lack of *Tdc2* neuronal signaling affects this coordination.

In addition to intraspecific matings, we used con- and heterospecific matings to show that inhibition of *Tdc2* neuron activity causes a higher P1 even when the first-male sperm is of lower compatibility to the female. This discovery motivates future studies to examine sperm handling dynamics and how sperm usage is coordinated with progeny production in this experimental setting to understand if the differential paternity proportion arises from the same mechanism as in conspecific matings.

In conclusion, our study demonstrates that the activity of octopaminergic/tyraminergic *Tdc2* neurons influences differential paternity proportions in doubly-mated females. *Tdc2* neural activity is required to mediate biased usage of the second male’s sperm, and allow females to produce mostly second-male progeny (low P1). Using an intersectional technique, we further show that this effect is largely mediated by *Tdc2* neurons innervating the female RT. On the other hand, over-activating *Tdc2* neurons does not further increase the second male’s paternity share, suggesting a ceiling effect. Future studies can investigate if *Tdc2* neuron activity also facilitates sperm preference based on male quality, and if other female neurons implicated in differential paternity outcome may exert their effects similarly through sperm preference or through other aspects of sperm handling. A better understanding of the molecular and physiological mechanisms underlying postcopulatory sexual selection may shed light on how female × male interactions drive evolution of reproductive traits.

## Acknowledgements

We are grateful to NIH R01 HD059060 (to AGC and MFW) for supporting this work. We thank members of the Clark and Wolfner labs for helpful discussions and suggestions, Dr. Carolina Rezával and Dr. Nilay Yapici for advice on *TNT* and *dTrpA1* experiments, Dr. John Belote, Dr. Stephen Goodwin, Dr. Scott Pitnick, Dr. Carolina Rezával, and Dr. Vanessa Ruta for fly strains, Dr. Ben Hopkins for the design of the mating plug ejection chambers, and Dr. Sofie Delbare and Orli Weiss for advice on heterospecific mating experiments.

